# Spherocytosis-related L1340P mutation in ankyrin affects its interactions with spectrin

**DOI:** 10.1101/2022.11.21.517333

**Authors:** Beata Machnicka, Aleksander Czogalla, Dżamila M. Bogusławska, Piotr Stasiak, Aleksander F. Sikorski

**Affiliations:** Department of Biotechnology, Institute of Biological Sciences, University of Zielona Góra, Prof. Z. Szafrana 1 St., PL 65-516 Zielona Góra, Poland; University of Wrocław, Faculty of Biotechnology, Department of Cytobiochemistry, F. Joliot-Curie 14a St., 50-383 Wrocław, Poland; Department of Anatomy and Histology, Collegium Medicum, University of Zielona Gora, Zyty 28 St., 65-046 Zielona Gora, Poland; Research and Development Centre, Regional Specialist Hospital, Kamieńskiego 73a St., 51-154 Wroclaw, Poland

**Keywords:** erythrocyte membrane skeleton, Ankyrin-1, Spectrin, Spectrin-binding domain, ankyrin-spectrin interaction, hereditary spherocytosis

## Abstract

Previously, we reported a new missense mutation in the *ANK1* gene correlated with the HS phenotype. This mutation, resulting in L1340P substitution (HGMD CM149731), likely leads to the changes in the conformation of the ankyrin ZZUD domain important for ankyrin binding to spectrin. In this report, we have shown the molecular and physiological effects of this mutation. First, we assessed the binding activity of human β-spectrin to the mutated ZZUDL1340P domain of ankyrin using two different experimental approaches – the study of association and dissociation responses of spectrin ankyrin binding domain and sedimentation assay. In addition, we demonstrated changes in morphology caused by the overexpressed ankyrin ZZUD domain in human cell models. Our results prove the key role of L1340 aa residue in the UPA domain for the correct alignment of the ZZUD domain of ankyrin, which results in binding the latter with spectrin within the erythrocyte membrane. Replacing the L1340 with a proline residue disrupts the spectrin binding activity of ankyrin.

## Introduction

The function of the erythrocyte membrane skeleton has been studied for years. Many interactions between membrane skeleton proteins and also membrane skeleton proteins with membrane bilayer containing integral membrane proteins via proteins and lipids were discovered due to identification of naturaly occuring mutants which are mostly related to haemolytic anaemias [1–3]. These mutants have become the subject of many studies to elucidate the regulatory mechanisms of spectrin-ankyrin interaction, which has significantly increased our knowledge of the structure and function of the erythrocyte membrane skeleton [4–10].

Hereditary spherocytosis (HS), reported worldwide and characterized by the presence of spherical-shaped erythrocytes (spherocytes) on the peripheral blood smear is the most common inherited anaemia (1 in 2000 births) in individuals of Caucasian ancestry [11,12]. This erythrocyte membranopathy refers to a group of heterogeneous anaemias that is most commonly associated with dominant inheritance, although non-dominant and recessive inheritance has also been described [13]. Many identified mutations have been found in genes encoding erythrocyte membrane proteins: ankyrin, anion exchanger 1 (AE1), β-spectrin, α-spectrin, or protein 4.2, which defects lead to reduced deformability and loss of membrane surface area [14–16]. These in turn lead to reduced deformability due to compromised integrity of the membrane skeleton [17]. The abnormal red blood cells (RBCs) are trapped and destroyed in the spleen, which is the main cause of hemolysis in HS [18]. HS is characterized by anaemia, jaundice (from haemolysis or biliary obstruction), splenomegaly with reticulocytosis, and increased osmotic fragility of RBCs, all with the background of a positive family history of the disease [19].

Hereditary spherocytosis is therefore caused by mutations in one of the genes encoding RBCs membrane proteins [11,20]. Overall, patients with dominant mutations are less affected than those with recessive defects. With the rapid development and wide application of gene diagnostic technologies, the detection rate of HS cases is increasing [10,21]. Most of the mutations arise in exons, but some occur in introns, suggesting that intronic mutations also play a significant role in the pathogenesis of hereditary spherocytosis [22]. The use of whole exome sequencing in recent times has made it possible to detect a particularly large number of new missense mutations, but most of them have not been described in terms of their functional consequences. Genetic screening in Human Genome Mutation Database (HGMD) has now identified several mutations in β-spectrin and ankyrin-R associated with HS (http://www.hgmd.cf.ac.uk). According to this database, to date 57 different β-spectrin mutations associated with HS phenotype, including 10 deletions, 6 insertions, 12 missense, 15 nonsense, and 11 splicing mutations, have been reported (accessed on 6 October 2022). Interestingly, it is relatively easy to characterize the possible mutations in human HS of the gene *SPTB*, which are located within the ankyrin binding domain: repeat 14 of β-spectrin - R1756X [23] and repeat 15 – A1884V [24]. This domain and its direct interactions with ankyrin-R are relatively well described [7,9].

Defects in ankyrin-R have been implicated in approximately half of all patients with hereditary spherocytosis [25,26]. Erythrocyte ankyrin contains an N-terminal membrane binding domain, a key spectrin binding domain (which includes a highly conserved ZZU tandem), and a C-terminal regulatory domain (which contains the death domain) [27–29]. According to HGMD and the literature, to date 93 different erythrocyte ankyrin-R mutations associated with HS phenotype, including 12 missense and 27 nonsense mutations, have been reported in human HS. Genetic screening has up to now identified 23 mutations in ZZUD tandem of ankyrin. According to HGMD and the literature, to date 93 different erythrocyte ankyrin-R mutations associated with HS phenotype, including 12 missense and 27 nonsense mutations, have been reported in human HS. Genetic screening has up to now identified 23 mutations in ZZUD tandem of ankyrin-R (according to HGMD), including 8 mutations located in the subdomain ZU5A, 7 in ZU5B, 3 in UPA (including R1252X, R1334X, and L1340P) and 5 in death domain (accessed on 6 October 2022).-R (according to HGMD), including 8 mutations located in the subdomain ZU5A, 7 in ZU5B, 3 in UPA (including R1252X, R1334X, and L1340P) and 5 in death domain (accessed on 6 October 2022). The region responsible for spectrin-binding comprise the ZU5A subdomain of erythrocyte ankyrin, which interacts with repeats 14-15 of β-spectrin [6,7].

In our previous study, we reported a new missense mutation in the *ANK1* gene correlated with the HS phenotype in a three-generation Polish family with autosomal dominant hereditary spherocytosis [30]. This mutation, resulting in L1340P (HGMD CM149731) substitution, leads probably to the changes in the ankyrin ZZUD domain interactions with spectrin. The leucine (L1340) is a conserved residue in human ankyrins (R, G, and B) [28] and is located in the UPA subdomain of the ankyrin R (NP_065209.2 ankyrin-1 isoform 1: L1340). Our previous modeling experiments showed that missense mutation L1340P affects the secondary structure of the ZZUD tandem (NP_065209.2 ankyrin-1 isoform 1: 913-1487 https://www.uniprot.org/uniprotkb/P16157/entry) [30]. Moreover, the substitution allows the rearrangement of several extra hydrogen bonds within the analyzed structure within the ZZUD tandem. As we have shown previously, two key phenylalanine residues, F913 and F916 are crucial for spectrin binding. Substitution of these residues led to a very high increase in the K_D_ value or abrogated binding completely [7]. The presented study aims to identify the molecular and physiological effects of the mutation in the *ANK1* gene encoding the erythrocyte membrane protein – ankyrin-R that leads to substitution L1340P. Our data provide new insight into the regulatory mechanisms of membrane skeletal proteins interaction pathways.

## Materials and methods

### Plasmids and bacterial strains

The plasmid pEGFP-C1, the cloning host *Escherichia coli* XL1-Blue strain was from Stratagene (USA), and the plasmid pRSETC and *E. coli* DH5α and *E. coli* BL21(DE3)pLysE strains were from Invitrogen (USA).

### Cloning, expression, and purification of His(6)GFP–AnkBD and GST-ZZUD fragments and its L1340P mutant

Ankyrin ZZUD supramodule was previously described as important for spectrin binding by ankyrin [30]. The 1746 bp length fragment of transcript variant 1 of the *ANK1* gene (NM_020476.3:2815-4560 bp) contained ankyrin ZZUD was synthesized at GenScript (International) employing *E*.*coli* codon optimization within pUC57 plasmid and subsequently cloned into pGEX-6P-1 high expression vector using Bam-HI and Xho-I restriction enzymes and standard ligation protocol. GST-ZZUD construct was amplified in XL1-Blue *E. coli* and positively screened plasmids were sequenced and transferred into expression BL21(DE3) pLysS *E. coli* strain using a standard protocol. AnkBD of spectrin 14-15 repeat was previously cloned into pRSETC expression vector proceeded by a sequence encoding Green Fluorescent Protein obtained from the pEGFP-C1 [7].

The site-directed mutagenesis was performed with QuickChange Site-Directed Mutagenesis Kit (Stratagene). The PCR products were stored in *E. coli* strain XL1-Blue. The resulting plasmids bearing mutated gene were sequenced to verify the mutations. Plasmid GST-ZZUDL1340P was then transferred into BL21(DE3) pLysS cells for overexpression of the proteins.

Overexpressed proteins: GST-ZZUD, GST-ZZUDL1340P, and His(6)GFP –AnkBD proteins were purified using Glutathione Sepharose 4B (GE Healthcare) or Talon® (Clontech) resins accordingly, as described previously [6]. The concentration of proteins was determined using a Cary 1E spectrophotometer employing extinction coefficient parameters determined for each protein [31]. Purity was also estimated from Coomassie stained SDS-PAGE on 10% gels (Figure S1).

### CD analysis

CD measurements were performed on a JASCO J-1500 CD spectrometer, using a thermostat-controlled cell with a 0.2 cm path length, from 20 to 70 °C in 5 °C increments, within the range of 205-260 nm. Ellipticity values from CD spectra were converted into molar residue ellipticity ([θ]M: in degrees cm^2^ dmol^−1^) values, as previous described [32].

### BLI method

Octet^®^ K2 2-channel System (ForteBio, Menlo Park, CA, USA) was used for binding studies. Experiments were performed in black 96 well plates (Nunc F96 MicroWellTMPlates, ThermoFisher Scientific). The total volume of each sample or buffer was 0.2 mL per well and shaking settings for every step were 1000 rpm. The test was performed at 30°C. Before each assay, Ni-NTA biosensors tips (ForteBio, Menlo Park, CA, USA) were pre wetted in 0.2 mL TRIS buffer (50 mM Tris–HCl, 150 mM NaCl, 2mMDTT, pH 7.5) for at least 10 min. Subsequently, the equilibration step with TRIS buffer was performed for 60 s, afterward, Ni-NTA biosensors tips were non-covalently loaded with His(6)GFP – AnkBD of spectrin for 600 s, followed by an additional equilibration step (250s) in TRIS buffer. The Association step of His(6)GFP –AnkBD with GST-ZZUD and GST-ZZUDL1340P batches in a concentration of ZZUD range 0 – 2,5 μM was carried out for 300 s. Dissociation was the last step and it lasted for 300 s. The response is measured as a shift in the interference pattern (in nm) and is proportional to the number of molecules bound to the surface of the biosensor. To analyze association and dissociation responses the curves were baseline corrected with the Octet Software (Version HT 11.1.0.25) and compared in terms of their amplitude for GST-ZZUD and GST-ZZUDL1340P.

### Sedimentation assay

The GST-tagged recombinant spectrin-binding domain of ankyrin (GST-ZZUD and GST-ZZUDL1340P) were adsorbed on a resin Glutathione sepharose 4B *via* GST tag. Saturated with a GST-tagged proteins resin were incubated with increasing concentrations range 0 – 2 nM of His(6)GFP–AnkBD of erythrocyte spectrin in the binding buffer (50 mMTris–HCl, 150 mM NaCl, 2mMDTT, pH 7.5) for 30 min at room temperature. The resin was then spun down at 1000×g for 5 min, resuspended in elution buffer with 75 mM reduced GSH, and centrifuged again. The amount of bound His(6)GFP– AnkBD in the supernatant was determined using a Varian Cary Eclipse Spectrofluorimeter. Control experiments were performed using the Glutathione sepharose 4B resin equilibrated in the test buffer and incubated with corresponding amounts of His(6)GFP–AnkBD. Bound values were obtained by subtracting control values from those for His(6)GFP–AnkBD bound to the resin saturated with GST-ZZUD or GST-ZZUDL1340P. The binding parameters were identified using one site - specific binding fitting (GraphPad Software Inc., San Diego, CA, USA).

### Resealed erythrocyte ghosts assay

Fresh blood samples were collected from healthy human volunteers (upon their informed, written consent) using anticoagulant (0.8% citric acid monohydrate, 2.2% trisodium citrate, 2.2% glucose). RBCs were washed three times with 10 mM Tris-HCl buffer (pH 7.4) containing 120 mM KCl. Intact cells were lysed in ice-cold lysis buffer containing 0.6 mM MgATP in 5 mM Tris-HCl, 5 mM KCl, 1 mM MgCl 2, pH 7.4, centrifuged (15 000 g, 15 min, 4oC) and washed in the same buffer until pale pink. The ghosts were resuspended in resealing buffer (150 mM KCl, 1.6 mM MgCl 2, 1 mM DTT, 0.6 mM MgATP, pH 7.4) and incubated for 60 min, at 37 °C in the presence or absence of recombinant ZZUD or its mutants (final concentration 1 mg/ml). After resealing, the suspension will be centrifuged (10 000 g, 5 min, 4°C), and pellets and supernatants will be collected for further analysis. Resealed ghosts were stained with lipophilic dye, Vybrant DiD (Molecular Probes), according to the manufacturer’s instructions and observed in the fluorescent microscope (Zeiss AXIO) as described previously [33,34].

### Cloning of ZZUD and its L1340P mutant domain into mEGPF –C1 and mRFP-C1 plasmids

An insert of ZZUD and ZZUDL1340P were cloned out from a positive pGEX-6P-1-ZZUD or pGEX-6P-1 - ZZUDL1340P with PCR based cloning procedure into a pEGFP-C1 (Addgene 54759) or mRFP-C1 (Addgene 54764) transfection vector coding for Green/Red Fluorescent Protein as a tag protein. Primers used for PCR cloning are shown in Table 1. Positive clones were screened as described above by using primers combination (Table 2) EGFP– ZZUD, EGFP-ZZUDL1340P, and mRFP-C1-ZZUD, mRFP-C1-ZZUDL1340P plasmids were purified using Plasmid Mini Kit (Eurex) and used for transfection.

**Table 1.**
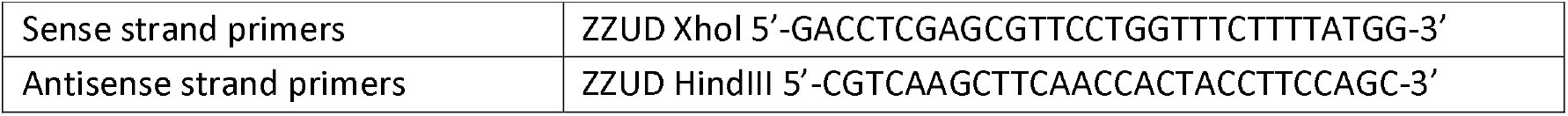
Primers used for PCR cloning

**Table 2.**
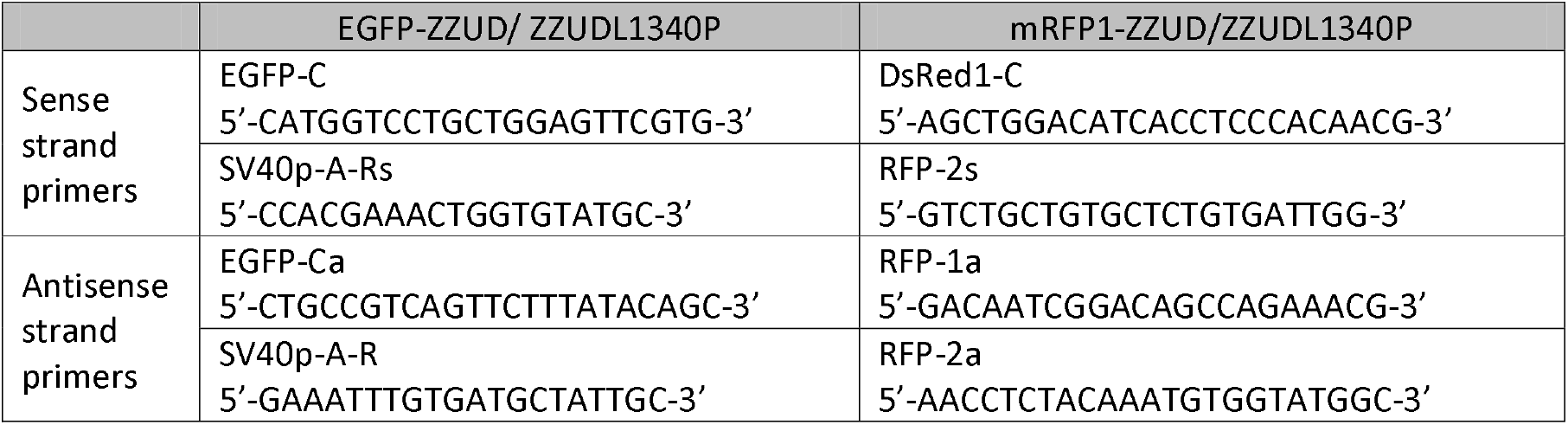
Primers used for sequencing

### HEL and K562 cells transfection

Before transfection HEL and K562 cells were grown directly on cover slips for 8 h in 6-well, 35 mm plates (Nunc™ Cell-Culture Treated Multidishes, ThermoFisher Scientific) in RPMI and IMEM medium respectively with 10% FCS and antibiotics (100 lg/ml streptomycin and 100 lg/ml penicillin), then transferred to medium without antibiotics. Cells were transfected with a Lipofectamine 2000 transfection reagent (Thermo Fisher) (μL) to DNA (μg) ratio of 3:1 mixture and cultured. Fluorescent microscopy assessed the changes in morphology resulting from overexpression of WT or mutated ZZUD fragments after 48 and 72 h (Zeiss AXIO).

## Results

### Binding studies of ankyrin and spectrin

The binding activity of β-spectrin is defined within the ZZUD domain (ZU5-ZU5-UPA-DD) of ankyrin. ZZUD arranged into the following tandem (NP_065209.2 ankyrin-1 isoform 1: 913-1487 https://www.uniprot.org/uniprotkb/P16157/entry): two tandem ZU5 domains (ZU5A and ZU5B) is composed of 156 and 147 residues, respectively), subdomain UPA (129 residues) and DD (termed as death domain, which contains 85 residues) (Figure S3). Repeats 14-15 of β-spectrin are responsible for interactions with the ZU5A subdomain of erythrocyte ankyrin [9]. The UPA domain could play a role in mediating the interactions between the ZU5B and DD domains [28,29]. To test the binding possibility of ankyrin ZZUD domain with spectrin, a fragment (NP_065209.2 ankyrin-1 isoform 1: 911-1492) of ankyrin was prepared. For the second construct, we introduced a point mutation within the ZZUD domain which mimics a natural mutation described previously [30]. A graphical representation of the used proteins construct is depicted in Figure 1.

**Figure 1.**
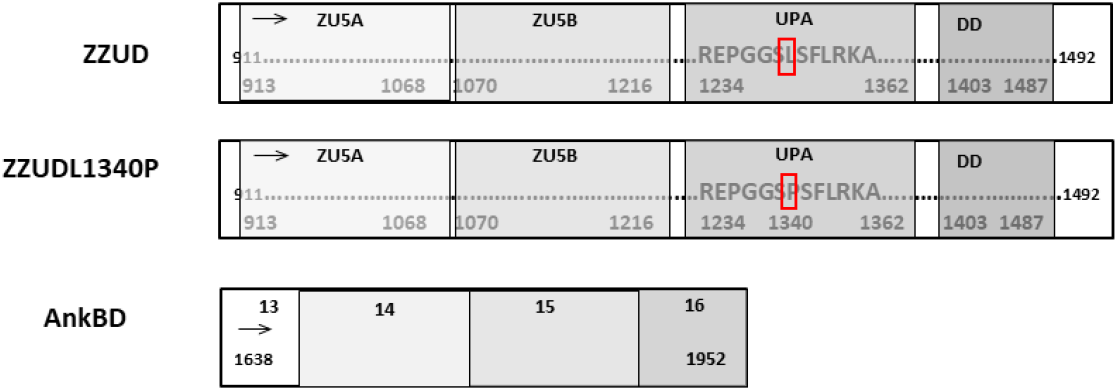
Schematic representation of the various ankyrin and spectrin fragments used in the assays concerning their length and domains.

Two independent approaches were used to measure and compare the binding activity of the spectrin to the ZZUD domain of ankyrin or its mutated version– ZZUDL1340P.

In a BLI assay, we measured the association and dissociation responses of ZZUD domain of ankyrin or its mutant to AnkBD of spectrin immobilized to the sensors using the two-channel Octet® K2 system (Figure 2A). We observed a significant decrease (approx. threefold) of binding response in the case of ankyrin fragment bearing L1340P mutation when compared to WT counterpart. This strongly suggests that ankyrin-spectrin interactions may be corrupted due to the L1340P mutation.

**Figure 2.**
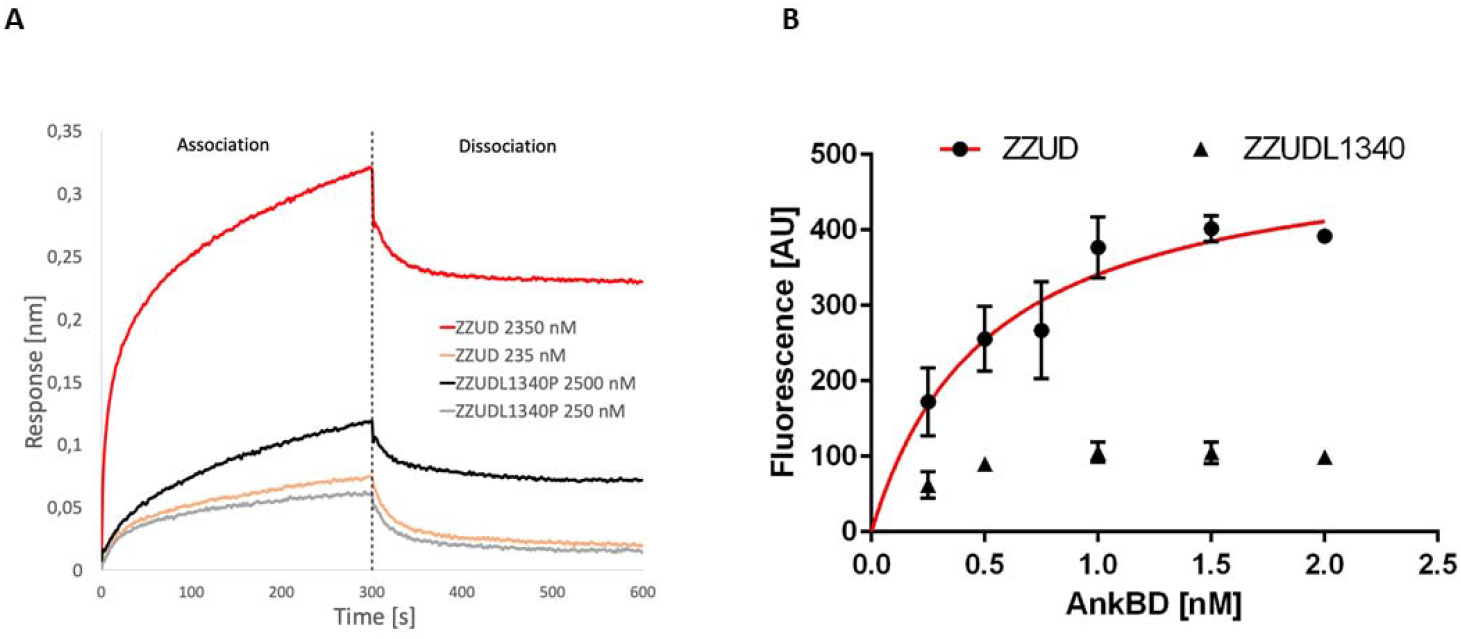
Quantification of binding between AnkBD and ZZUD or its mutated form. (A) Binding of ZZUD or ZZUDL1340P mutant to BLI sensor with AnkBD. (B) Saturation binding curve of ZZUD and AnkBD obtained via sedimentation assay.

To get more quantitative data on the influence of L1340P mutation on spectrin-ankyrin interactions in parallel to BLI we employed sedimentation assay (Figure 2B), using GFP labelled AnkBD of spectrin and GST-ZZUD domain of ankyrin. Having saturation binding curves, we managed to calculate the equilibrium dissociation constant. The value of the equilibrium dissociation constant K_D_=0.519nM indicates that AnkBD of spectrin and ZZUD domain of ankyrin complexes of high affinity are formed and is similar to earlier values reported by others [4,35] or by us [7]. Whereas for the mutated form of ankyrin-derived fragment (ZZUDL1340P), it was not possible to calculate K_D_ due to the very low fluorescence values obtained (Bmax approx. 5 times lower than in case of wild type domain). These results showed that the presence of the L1340P mutation in the ZZUD domain significantly disrupts the ability to form a high affinity complex. Moreover, the results obtained from the ZZUDL1340P fragment further confirm the importance of leucine residue in the proper binding of spectrin with ankyrin, as we suggested previously [30].

Circular dichroism (CD) analysis confirmed that the loss of binding was not caused by the instability of the mutant. The actual T_m_ of the mutant was higher than that for the WT ZZUD domain (Figure S2B). Both protein domains are characterized by a high level of stability and undergo similar folding. AnkBD of spectrin was also confirmed to be properly folded (Figure S2). However, the recorded CD spectra showed that the L1340P mutation results in some very minor changes in the secondary structure (Figure S2). Slight changes in one of the β-sheets were also visible in the model which we previously generated [30]. The modelling experiments performed there showed that the substitution L1340P affects structural changes in the beta strand which is longer and the orientation of F913 residue is changed.

### Effect of mutations on the morphology of resealed erythrocyte ghosts

The effect of encapsulation of the WT or the mutated ZZUD domain of ankyrin in the erythrocyte ghost on their morphology was followed. For this purpose, red blood cells were prepared, loaded with the purified wild-type ZZUD domain and its L1340P mutant, and then resealed ghosts were stained with DiD dye. It was observed that the ghosts of erythrocytes loaded with the non-mutated ZZUD domain were characteristically deformed, while those loaded with the purified mutated ZZUD L1340P domain did not change their shape (Figure 3). These results indicate the lack of ability of ZZUDL1340P to compete for binding erythrocyte ghost spectrin with native ankyrin residing within membrane skeleton responsible for shape maintenance. No changes in erythrocyte ghost morphology in the presence of the mutant form suggests the importance of the mutant L1340P residue in maintaining the mechanical properties of the membrane and the shape of the erythrocytes.

**Figure 3.**
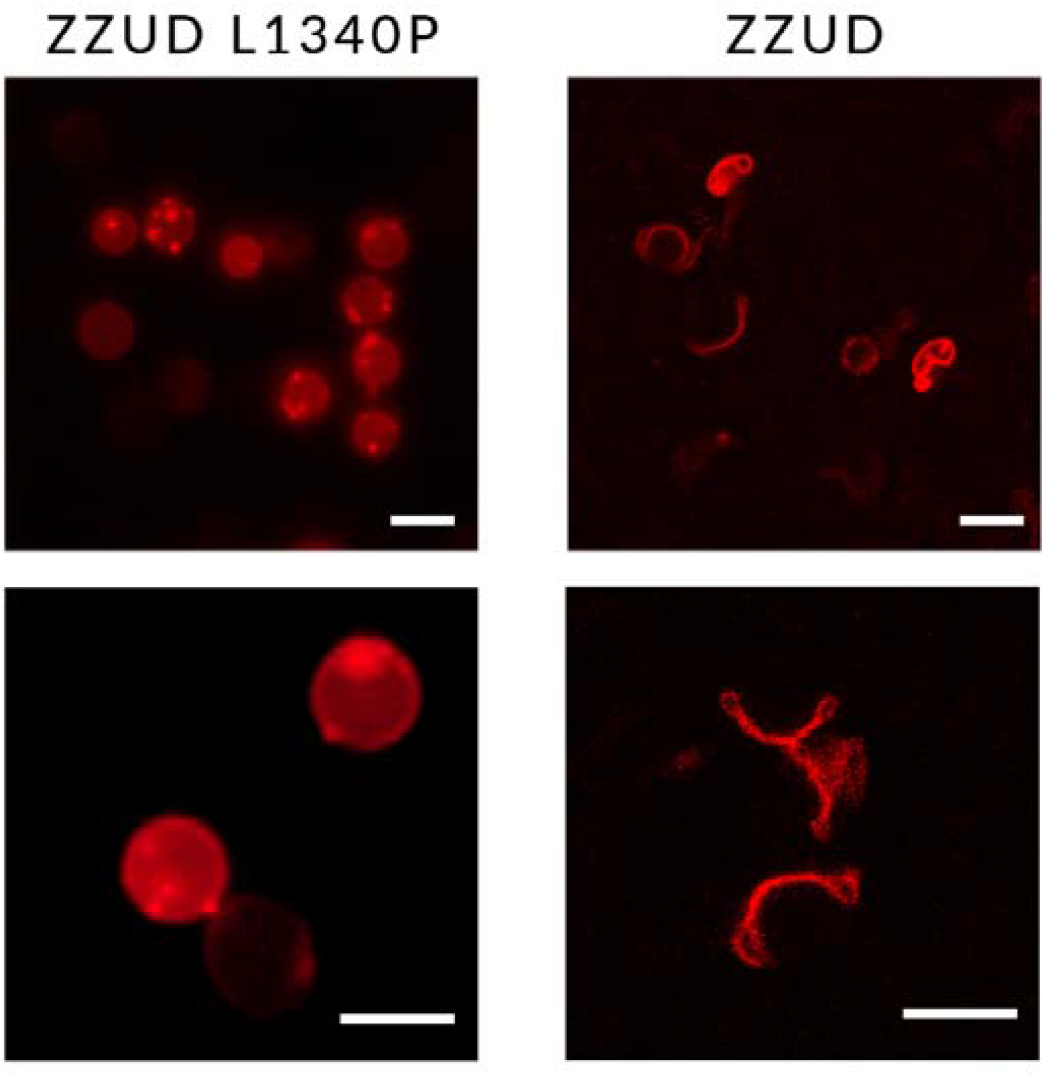
Effect of the ZZUD domain and its mutant form on the morphology of the resealed erythrocyte ghosts. The ghosts of erythrocytes loaded with the non-mutated ZZUD domain were characteristically deformed (right panel), while those loaded with the purified mutated ZZUD domain remained spherical (left panel).

### Effect mutations on the shape of transiently transfected HEL and K562 cells

To assess the effect of the overexpression of the spectrin binding domain of ankyrin ZZUD on the living cells *in vitro*, HEL, and K562 cells were transiently transfected with vectors carrying inserts encoding protein fragments fused with GFP or RFP as described by Bok et al. [36]. The changes in morphology resulting from overexpression of ZZUD or mutated ZZUDL1340P fragments of ankyrin were observed in fluorescent microscopy.

For comparison, the constructs carrying spectrin–binding domain or its L1340P mutant were overexpressed in erythroblast cells line HEL. The results are presented in Figure 4A. HEL cells transfected with the vector encoding fluorescent protein only were used as a control. The first effect of overexpression was observed 48h after transfection. RFP and RFP-ZZUDL1340P overexpression did not change the morphology of the cells. In contrast, overexpression of erythroid ankyrin’s ZZUD causes granulation along the membrane surface. The effect is more visible 3 days after transfection. 96h after transfection, the cells overexpressed ZZUD-RFP were mostly damaged. Four day after transfection, almost all ZZUDL1340P transfected cells were dead in contrast with cells overexpressed mutated fragment of ankyrin. Cells transfected with control plasmid and a plasmid carrying insert encoding mutated ZZUDL1340P domain remained over all time of experiment still alive without any changes in cell shape.

**Figure 4.**
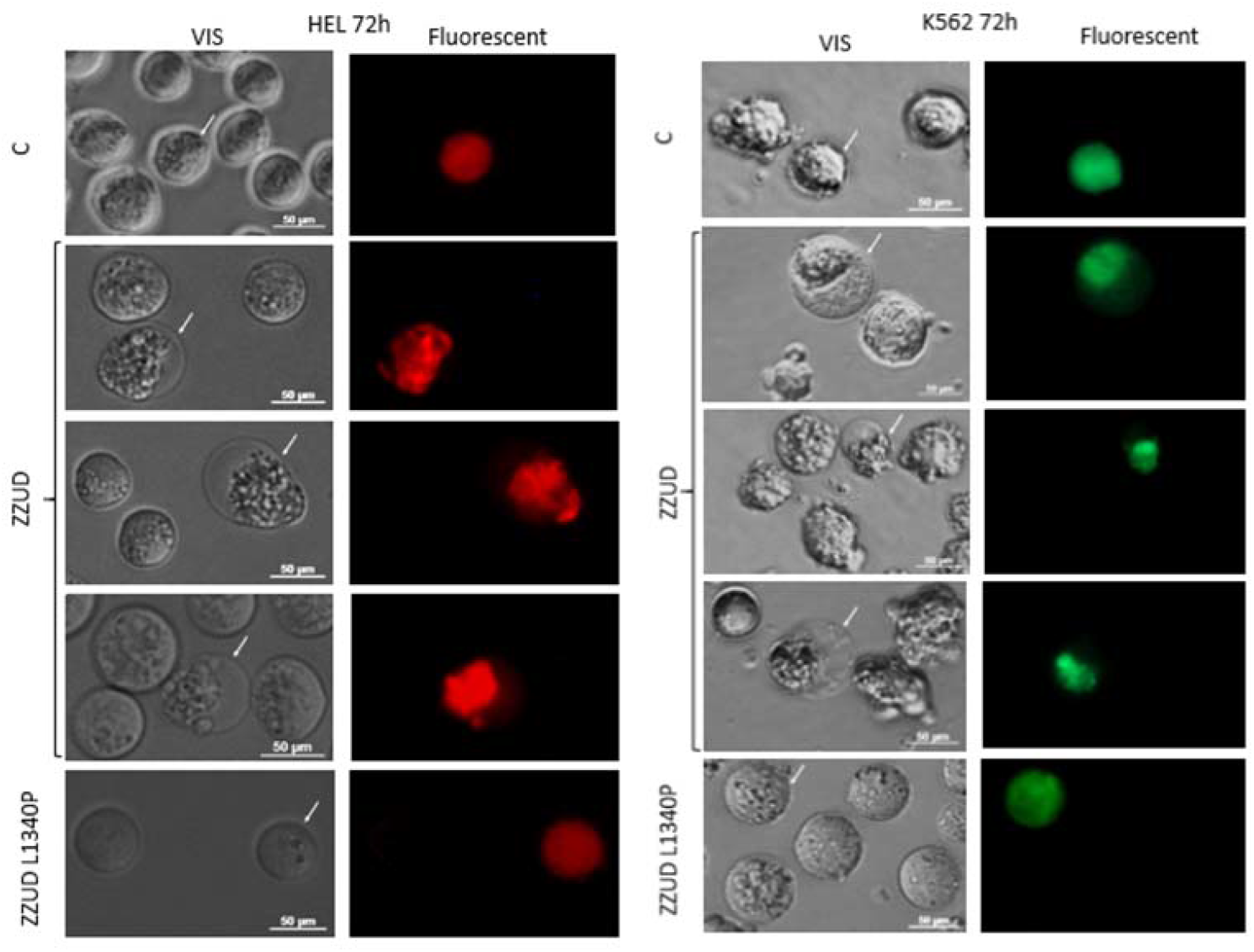
HEL and K562 cells were transfected with EGFP/RFP ZZUD and EGFP/ RFP ZZUDL1340P plasmids 72 hours after transfection. The cells continue to have a circular outline, but the ankyrin-drived fragments accumulates at a certain region of the cell leading to its punctate occurrence - such appearance is not present in the wild-type/control cells or ZZUD L1340P transfected ones.

The similar observations were provided using lymphoblast cell line K562 (Figure 4B). The overexpression of the ZZUD domain causes changes in cell morphology 48 and 72h after transfection. The observed cells had an increasing aggregation pattern.

These results indicate that wild type of ZZUD domain when transiently overexpressed in living cells induced aggregation of the spectrin based membrane skeleton leading to its punctate appearance while no effect was observed for ZZUD L1340P mutant.

## Discussion

In hereditary spherocytosis (HS), the inappropriate binding of spectrin to ankyrin is highly correlated with severity of the disease [15,28]. The region responsible for proper spectrin-binding is the ZU5A subdomain of erythrocyte ankyrin, which interacts with repeats 14-15 of β-spectrin [4,5].

The structural studies indicate that the second ZU5B domain is involved in the formation of an auto-regulating supramodule ZZUD of ankyrin [29]. Based on the analysis of eight ankyrin-B syndrome-causing mutations (including two located in UPA) Wang et al. suggested that the ankyrin-B ZZUD tandem is likely to bind to other proteins, thereby regulating its N-terminal ankyrin repeats-mediated membrane binding. Other missense mutation W1185R [28] in ankyrin-R, which is located in the ZU5B domain and not directly involved in spectrin binding, can cause hereditary spherocytosis. This further indicates that this ZU5 domain plays important functional role within ankyrins.

The main factor in the correct binding of ankyrin and spectrin is the presence of complementary charged surfaces on these proteins that match each other [7]. Pathological lesions in HS result mainly from the loss of spectrin and/or ankyrin that is normally responsible for anchoring the membrane skeleton in the lipid bilayer. The current knowledge is constantly updated with new mutations resulting in the HS phenotype [10,37–40].

In our previous study, we described a new missense mutation resulting in L1340P (HGMD CM149731) substitution in the *ANK1* gene correlated with the HS phenotype [30]. The aim of this project was to perform *in vitro* experimental analyzes using purified ankyrin and spectrin protein domains previously described as being involved in interactions with each other and to determine the impact of the L1340P mutation found in ankyrin on the interaction between these proteins. For the purposes of the study the stable construct of ZZUD fragments of ankyrin with a mutation which mimics a natural mutation L1340P was prepared. The applied strategy allowed us to purify bacterially expressed and properly folded proteins. CD spectra suggest that both fragments correspond to the native protein. The binding activity of human β-spectrin with mutated ZZUDL1340P domain of ankyrin was assessed using two different experimental approaches – sedimentation assay and study of association and dissociation responses in respect to spectrin ankyrin binding domain via biolayer interferometry. Obtained *K*_*D*_ values of ZZUD domain – AnkBD of spectrin complexes highly corresponded to values obtained previously for interaction between spectrin and ankyrin by different teams [6,27]. The binding affinity of spectrin to mutated ZZUDL1340P domain decreased dramatically when compared to the control wild type ZZUD domain.

In our next research, the lack of ability of the mutated fragment of ZZUD to compete for binding with erythrocyte ghost spectrin was demonstrated. No changes in erythrocyte ghost morphology in the presence of the mutant form suggests that the mutant L1340P does not compete for spectrin with native ankyrin. Thus, aa residue at 1340 position is critical in maintaining the mechanical properties of the membrane and the shape of the erythrocytes. Moreover, in cellular models, we demonstrated changes in morphology cells overexpressed the ZZUD domain of ankyrin. In the transfected erythroblast cells line, HEL and lymphoblast cells K562 expressing GFP/RFP-ZZUD proteins causes large granulation along the membrane surface of both cell types. Probably spectrin binding fragment of ankyrin is competitive for endogenous spectrin in ankyrin binding activity. It should be noted that with time number of cells overexpressed the ZZUD domain dramatically decreased. This was probably caused by enhanced mortality of these cells due to the presence of more ankyrin fragments which destabilized the membrane. Similar phenotype we observed previously in HeLa cells overexpressed ankyrin binding domain of spectrin [6].

Our *in vitro* and *in cellula* results prove the key role of L1340 in the UPA domain for the proper function of the ZZUD domain resulting in the ability to form a bond between spectrin and ankyrin in the erythrocyte membrane. Replacing L1340 with a proline residue disrupts the activity of ankyrin to bind spectrin, which is correlated with the HS phenotype of patients with autosomal dominant hereditary spherocytosis.

## Supporting information

MachnickaetalSupplfile

## Acknowledgement

This work was supported by grant from the National Science Centre, Poland Project No. 2015/19/B/NZ5/03469.

## References

[1] J. Narla, N. Mohandas, Red cell membrane disorders, Int J Lab Hem. 39 (2017) 47–52. https://doi.org/10.1111/ijlh.12657.

[2] J. Pugi, L.J. Drury, J.C. Langer, S. Butchart, M. Fantauzzi, J. Baker, V.S. Blanchette, M.A. Kirby-Allen, M. Carcao, Genotype/Phenotype Correlations in 103 Children from 87 Families with Hereditary Spherocytosis, Blood. 128 (2016) 2432–2432. https://doi.org/10.1182/blood.V128.22.2432.2432.

[3] N. Mohandas, Inherited hemolytic anemia: a possessive beginner’s guide, Hematology. 2018 (2018) 377–381. https://doi.org/10.1182/asheducation-2018.1.377.

[4] J.J. Ipsaro, L. Huang, A. Mondragón, Structures of the spectrin-ankyrin interaction binding domains, Blood. 113 (2009) 5385–5393. https://doi.org/10.1182/blood-2008-10-184358.

[5] J.J. Ipsaro, A. Mondragón, Structural basis for spectrin recognition by ankyrin, Blood. 115 (2010) 4093–4101. https://doi.org/10.1182/blood-2009-11-255604.

[6] A. Kolondra, M. Grzybek, A. Chorzalska, A.F. Sikorski, The 22.5kDa spectrin-binding domain of ankyrinR binds spectrin with high affinity and changes the spectrin distribution in cells in vivo, Protein Expression and Purification. 60 (2008) 157–164. https://doi.org/10.1016/j.pep.2008.04.002.

[7] A. Kolondra, M. Lenoir, M. Wolny, A. Czogalla, M. Overduin, A.F. Sikorski, M. Grzybek, The role of hydrophobic interactions in ankyrin–spectrin complex formation, Biochimica et Biophysica Acta (BBA) -Biomembranes. 1798 (2010) 2084–2089. https://doi.org/10.1016/j.bbamem.2010.07.024.

[8] P.J. La-Borde, P.R. Stabach, I. Simonović, J.S. Morrow, M. Simonović, Ankyrin recognizes both surface character and shape of the 14–15 di-repeat of β-spectrin, Biochemical and Biophysical Research Communications. 392 (2010) 490–494. https://doi.org/10.1016/j.bbrc.2010.01.046.

[9] A. Czogalla, A.F. Sikorski, Do we already know how spectrin attracts ankyrin?, Cell. Mol. Life Sci. 67 (2010) 2679–2683. https://doi.org/10.1007/s00018-010-0371-1.

[10] D.M. Bogusławska, M. Skulski, B. Machnicka, S. Potoczek, S. Kraszewski, K. Kuliczkowski, A.F. Sikorski, Identification of a Novel Mutation of β-Spectrin in Hereditary Spherocytosis Using Whole Exome Sequencing, IJMS. 22 (2021) 11007. https://doi.org/10.3390/ijms222011007.

[11] J. Delaunay, The molecular basis of hereditary red cell membrane disorders, Blood Rev. 21 (2007) 1–20. https://doi.org/10.1016/j.blre.2006.03.005.

[12] L. Da Costa, J. Galimand, O. Fenneteau, N. Mohandas, Hereditary spherocytosis, elliptocytosis, and other red cell membrane disorders, Blood Reviews. 27 (2013) 167–178. https://doi.org/10.1016/j.blre.2013.04.003.

[13] P. Agre, A. Asimos, J.F. Casella, C. McMillan, Inheritance Pattern and Clinical Response to Splenectomy as a Reflection of Erythrocyte Spectrin Deficiency in Hereditary Spherocytosis, N Engl J Med. 315 (1986) 1579–1583. https://doi.org/10.1056/NEJM198612183152504.

[14] D.M. Bogusławska, E. Heger, K. Baldy-Chudzik, M. Zagulski, M. Maciejewska, A. Likwiarz, A.F. Sikorski, (AC)n microsatellite polymorphism and 14-nucleotide deletion in exon 42 ankyrin-1 gene in several families with hereditary spherocytosis in a population of South-Western Poland, Ann Hematol. 85 (2006) 337–339. https://doi.org/10.1007/s00277-006-0083-7.

[15] S. Eber, S.E. Lux, Hereditary spherocytosis—defects in proteins that connect the membrane skeleton to the lipid bilayer, Seminars in Hematology. 41 (2004) 118–141. https://doi.org/10.1053/j.seminhematol.2004.01.002.

[16] G. Paździor, M. Langner, A. Chmura, D. Bogusławska, E. Heger, A. Chorzalska, A.F. Sikorski, The kinetics of haemolysis of spherocytic erythrocytes, Cell Mol Biol Lett. 8 (2003) 639–648.

[17] D.O. Demiralp, S. Peker, B. Turgut, N. Akar, Comprehensive identification of erythrocyte membrane protein deficiency by 2D gel electrophoresis based proteomic analysis in hereditary elliptocytosis and spherocytosis, Prot. Clin. Appl. 6 (2012) 403–411. https://doi.org/10.1002/prca.201200010.

[18] R. Reliene, M. Mariani, A. Zanella, W.H. Reinhart, M.L. Ribeiro, E.M. del Giudice, S. Perrotta, A. Iolascon, S. Eber, H.U. Lutz, Splenectomy prolongs in vivo survival of erythrocytes differently in spectrin/ankyrin- and band 3-deficient hereditary spherocytosis, Blood. 100 (2002) 2208–2215.

[19] M.-J. King, A. Zanella, Hereditary red cell membrane disorders and laboratory diagnostic testing, Int J Lab Hematol. 35 (2013) 237–243. https://doi.org/10.1111/ijlh.12070.

[20] W.T. Tse, S.E. Lux, Red blood cell membrane disorders, Br J Haematol. 104 (1999) 2–13. https://doi.org/10.1111/j.1365-2141.1999.01130.x.

[21] H. Li, D. Papageorgiou, H.-Y. Chang, L. Lu, J. Yang, Y. Deng, Synergistic Integration of Laboratory and Numerical Approaches in Studies of the Biomechanics of Diseased Red Blood Cells, Biosensors. 8 (2018) 76. https://doi.org/10.3390/bios8030076.

[22] B.-J. He, L. Liao, Z.-F. Deng, Y.-F. Tao, Y.-C. Xu, F.-Q. Lin, Molecular Genetic Mechanisms of Hereditary Spherocytosis: Current Perspectives, Acta Haematol. 139 (2018) 60–66. https://doi.org/10.1159/000486229.

[23] M. Maciag, D. Płochocka, A. Adamowicz-Salach, B. Burzyńska, Novel beta-spectrin mutations in hereditary spherocytosis associated with decreased levels of mRNA, Br J Haematol. 146 (2009) 326–332. https://doi.org/10.1111/j.1365-2141.2009.07759.x.

[24] P.G. Gallagher, B.G. Forget, Hematologically Important Mutations: Spectrin and Ankyrin Variants in Hereditary Spherocytosis, Blood Cells, Molecules, and Diseases. 24 (1998) 539–543. https://doi.org/10.1006/bcmd.1998.0217.

[25] P.G. Gallagher, Hematologically important mutations: ankyrin variants in hereditary spherocytosis, Blood Cells Mol Dis. 35 (2005) 345–347. https://doi.org/10.1016/j.bcmd.2005.08.008.

[26] A.M. Agarwal, Ankyrin Mutations in Hereditary Spherocytosis, Acta Haematol. 141 (2019) 63–64. https://doi.org/10.1159/000495339.

[27] A. Hryniewicz-Jankowska, A. Czogalla, E. Bok, A.F. Sikorsk, Ankyrins, multifunctional proteins involved in many cellular pathways, Folia Histochem Cytobiol. 40 (2002) 239–249.

[28] C. Wang, C. Yu, F. Ye, Z. Wei, M. Zhang, Structure of the ZU5-ZU5-UPA-DD tandem of ankyrin-B reveals interaction surfaces necessary for ankyrin function, Proceedings of the National Academy of Sciences. 109 (2012) 4822–4827. https://doi.org/10.1073/pnas.1200613109.

[29] M. Yasunaga, J.J. Ipsaro, A. Mondragón, Structurally Similar but Functionally Diverse ZU5 Domains in Human Erythrocyte Ankyrin, Journal of Molecular Biology. 417 (2012) 336–350. https://doi.org/10.1016/j.jmb.2012.01.041.

[30] D.M. Bogusławska, E. Heger, M. Listowski, D. Wasiński, K. Kuliczkowski, B. Machnicka, A.F. Sikorski, A novel L1340P mutation in the ANK1 gene is associated with hereditary spherocytosis?, Br. J. Haematol. 167 (2014) 269–271. https://doi.org/10.1111/bjh.12960.

[31] E. Gasteiger, C. Hoogland, A. Gattiker, S. Duvaud, M.R. Wilkins, R.D. Appel, A. Bairoch, Protein Identification and Analysis Tools on the ExPASy Server, in: J.M. Walker (Ed.), The Proteomics Protocols Handbook, Humana Press, Totowa, NJ, 2005: pp. 571–607. https://doi.org/10.1385/1-59259-890-0:571.

[32] E. Bilkova, R. Pleskot, S. Rissanen, S. Sun, A. Czogalla, L. Cwiklik, T. Róg, I. Vattulainen, P.S. Cremer, P. Jungwirth, Ü. Coskun, Calcium Directly Regulates Phosphatidylinositol 4,5-Bisphosphate Headgroup Conformation and Recognition, J. Am. Chem. Soc. 139 (2017) 4019–4024. https://doi.org/10.1021/jacs.6b11760.

[33] A. Chorzalska, A. Łach, T. Borowik, M. Wolny, A. Hryniewicz-Jankowska, A. Kolondra, M. Langner, A. Sikorski, The effect of the lipid-binding site of the ankyrin-binding domain of erythroid β-spectrin on the properties of natural membranes and skeletal structures, Cellular and Molecular Biology Letters. 15 (2010). https://doi.org/10.2478/s11658-010-0012-6.

[34] M. Wolny, M. Grzybek, E. Bok, A. Chorzalska, M. Lenoir, A. Czogalla, K. Adamczyk, A. Kolondra, W. Diakowski, M. Overduin, A.F. Sikorski, Key Amino Acid Residues of Ankyrin-Sensitive Phosphatidylethanolamine/Phosphatidylcholine-Lipid Binding Site of βI-Spectrin, PLoS ONE. 6 (2011) e21538. https://doi.org/10.1371/journal.pone.0021538.

[35] J.J. Ipsaro, L. Huang, L. Gutierrez, R.I. MacDonald, Molecular Epitopes of the Ankyrin−Spectrin Interaction †, Biochemistry. 47 (2008) 7452–7464. https://doi.org/10.1021/bi702525z.

[36] E. Bok, E. Plazuk, A. Hryniewiczjankowska, A. Chorzalska, A. Szmaj, P. Dubielecka, K. Stebelska, W. Diakowski, M. Lisowski, M. Langner, Lipid-binding role of βII-spectrin ankyrin-binding domain, Cell Biology International. 31 (2007) 1482–1494. https://doi.org/10.1016/j.cellbi.2007.06.014.

[37] S. Chai, R. Jiao, X. Sun, P. Fu, Q. Zhao, M. Sang, Novel nonsense mutation p. Gln264Ter in the ANK1 confirms causative role for hereditary spherocytosis: a case report, BMC Med Genet. 21 (2020) 223. https://doi.org/10.1186/s12881-020-01161-4.

[38] L. Hao, S. Li, D. Ma, S. Chen, B. Zhang, D. Xiao, J. Zhang, N. Jiang, S. Jiang, J. Ma, Two novel ANK1 loss-of-function mutations in Chinese families with hereditary spherocytosis, J Cell Mol Med. 23 (2019) 4454–4463. https://doi.org/10.1111/jcmm.14343.

[39] R. Russo, I. Andolfo, F. Manna, A. Gambale, R. Marra, B.E. Rosato, P. Caforio, V. Pinto, P. Pignataro, K. Radhakrishnan, S. Unal, G. Tomaiuolo, G.L. Forni, A. Iolascon, Multi-gene panel testing improves diagnosis and management of patients with hereditary anemias, Am J Hematol. 93 (2018) 672–682. https://doi.org/10.1002/ajh.25058.

[40] D.M. Bogusławska, E. Heger, B. Machnicka, M. Skulski, K. Kuliczkowski, A.F. Sikorski, A new frameshift mutation of the β-spectrin gene associated with hereditary spherocytosis, Ann Hematol. 96 (2017) 163–165. https://doi.org/10.1007/s00277-016-2838-0.

