## Supplementary material for "Spherocytosis-related L1340P mutation in ankyrin affects its interactions with spectrin": MachnickaetalSupplfile

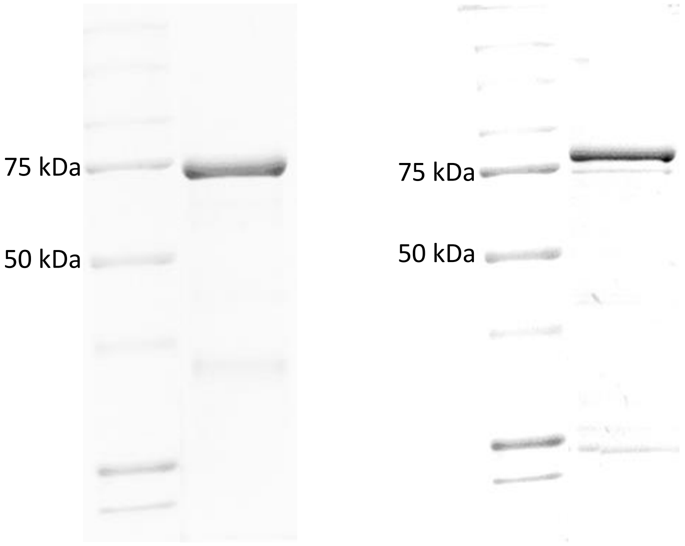


A B

Figure S1. Representative SDS-PAGE of purified His(6)GFP-AnkBD (A) and ZZUD-GST (B). Samples were run in 10% gel and stained with Coomassie Blue. Standard was Precision Plus Unstained Protein Standard (Bio-Rad).

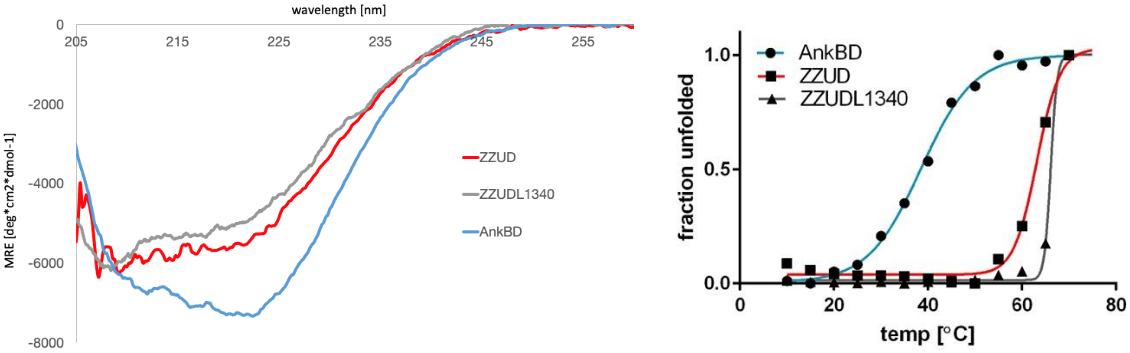


A B

Figure S2. Representative circular dichroism spectra (A) measured at 20 °C and melting curves (B) of His(6)GFP-AnkBD (blue), ZZUD-GST (red) and ZZUD L1340-GST (gray).

**ZU5 1**
>sp|P16157|ANK1_HUMAN|913-1068 OS=Homo sapiens OX=9606 GN=ANK1 PE=1 SV=3
FLVSFMVDAR GGSMRGSRHN GLRVVIPPRT CAAPTRITCR LVKPQKLSTP PPLAEEEGLA
SRIIALGPTG AQFLSPVIVE IPHFASHGRG DRELVVLRSE NGSVWKEHRS RYGESYLDQI
LNGMDEELGS LEELEKKRVC RIITTDFPLY FVIMSR
**ZU5 2**
>sp|P16157|ANK1_HUMAN|1070-1216 OS=Homo sapiens OX=9606 GN=ANK1 PE=1 SV=3
CQDYDTIGPE GGSLKSKLVP LVQATFPENA VTKRVKLALQ AQPVPDELVT KLLGNQATFS
PIVTVEPRRR KFHRPIGLRI PLPPSWTDNP RDSGEGDTTS LRLLCSVIGG TDQAQWEDIT
GTTKLVYANE CANFTTNVSA RFWLSDC
**UPA**
>sp|P16157|ANK1_HUMAN|1234-1362 OS=Homo sapiens OX=9606 GN=ANK1 PE=1 SV=3
TAVPYMAKFV IFAKMNDPRE GRLRCYCMTD DKVDKTLEQH ENFVEVARSR DIEVLEGMSL
FAELSGNLVP VKKAAQQRSF HFQSFRENRL AMPVKVRDSS REPGGSLSFL RKAMKYEDTQ
HILCHLNIT
**Death**
>sp|P16157|ANK1_HUMAN|1403-1487 OS=Homo sapiens OX=9606 GN=ANK1 PE=1 SV=3
AEMKMAVISE HLGLSWAELA RELQFSVEDI NRIRVENPNS LLEQSVALLN LWVIREGQNA
NMENLYTALQ SIDRGEIVNM LEGSG

Figure S3. The sequence of ZZUD supramodule according UNIPROT database
https://www.uniprot.org/uniprotkb/P16157/entry
